# A statistical framework for disease classification with scRNA-Seq Data

**DOI:** 10.64898/2026.08.21.746294

**Authors:** Zhiwei Xiao, William Torous, Jeffrey Cheng, Raymond Cho, Elizabeth Purdom

## Abstract

**Motivation:** Bulk RNA-sequencing based disease classification obscures cell-type specific signals by aggregating gene expression across heterogeneous tissues. Although single-cell RNA-seq tackles this limitation, summarizing and deriving patient-level predictors while retaining biological interpretability remains challenging. Standard sparse methods, such as lasso, often select arbitrary scattered gene sets without leveraging the underlying cell type structures revealed by single-cell data.

**Results:** We introduce a two-stage statistical framework for interpretable patient-level disease classification from single-cell data. We first construct a gene-by-cell-type pseudobulk matrix that summarize single-cell expression for each patient. We then fit a multinomial logistic regression model with sparse group lasso penalty, inducing sparsity at both the cell type and gene levels. Across datasets of systemic lupus erythematosus, COVID-19, and colorectal cancer, our framework either matched or outperformed lasso and random forest baselines. Importantly, our models recovered biologically coherent, cell-type specific gene signatures consistent with known disease mechanisms, demonstrating improved interpretability without sacrificing predictive accuracy.

**Availability:** The `scSGL` R package implementing the Sparse Group Lasso classification framework described in this paper is available at https://github.com/zhiweixiao/scSGL (version 0.99.1). Code to reproduce the actual cross-validation, model fitting, and prediction analyses on the three datasets reported here is available at https://github.com/zhiweixiao/scSGL-manuscript.

## 1 Introduction

### 1.1 Background

Predicting disease or phenotypes from gene expression data has been a long-standing strategy, particularly with sparse prediction algorithms like lasso [1] that provide interpretable gene signatures. However, these standard bulk measurements of mRNA average gene expression across heterogeneous cell populations, potentially masking critical cell-type-specific signatures that govern disease pathways [2]. Disease-informative cells present in small numbers can become undetectable when their signals are averaged with other abundant, healthy or noninformative cells.

Single-cell technologies can overcome the inherent limitations of traditional bulk approaches [3]. At the individual cellular level, these technologies can reveal cellular heterogeneity within seemingly uniform tissue samples, identify rare disease-driving cell populations, and enable the discovery of cell-type-specific biomarkers and therapeutic targets that bulk methods systematically miss [4].

### 1.2 Motivation

However, a fundamental challenge in utilizing whole-sample single-cell data lies in effectively extracting information from over thousands of individual cells per patient into meaningful predictors for classification models. From a statistical perspective, the objective is patient-level disease classification rather than individual cell classification. Classifying individual cells would therefore be inappropriate since disease diagnoses apply to patients, not cells, and not all cells from a diseased patient would necessarily show pathological behavior. Therefore, the challenge becomes how to aggregate cellular information while preserving the disease-discriminating patterns that emerge from specific cell types.

Furthermore, while sparse models such as lasso are often appealing for high dimensional genomic data due to their simplicity and interpretability, the relevant features of single-cell applications differ fundamentally from those of bulk data. In particular, meaningful predictors must capture both the expression of individual genes as well as the cellular structure underlying single-cell measurements.

### 1.3 Our solution

To address these challenges, we propose a two-stage prediction framework that first represents single-cell data as interpretable patient-level summaries in pseudobulk formats, and then applies a structured regularization technique for cell-type-specific predictor discovery.

In the first stage, we adopt a pseudobulk representation strategy [5] that summarizes the single-cell information into patient-level cell-type-specific expression profiles. This representation provides computational efficiency and biological interpretability while preserving the cell-type resolution essential for understanding disease mechanisms, and it scales naturally to large cohorts.

However, this pseudobulk transformation remains high-dimensional and requires appropriate regularization for patient-level prediction. In this setting, standard lasso regularization (L1) is suboptimal: it treats all features independently and therefore tends to select scattered individual genes across many different cell types without reference to their cellular origin [6]. Such fragmented selection obscures the underlying cellular structure of disease and complicates further biological interpretation. Moreover, genes expressed within the same cell type often exhibit strong correlation due to shared regulatory mechanisms, and standard lasso’s tendency to arbitrarily select among correlated features can further reduce predictive algorithms’ interpretability and stability [7].

To overcome this, we instead use the sparse group lasso (SGL) penalty of [8] within a multinomial logistic regression framework. The SGL penalty inherently aligns with the hierarchical structure of our pseudobulk data by inducing sparsity at two levels simultaneously: sparsity of cell-types and sparsity of genes within cell-types. As a result, the model provides clear biological interpretation about which cellular populations drive disease classification, and within such populations, which gene signatures are the most predictive.

Compared to more sophisticated approaches such as deep learning models that can achieve high predictive accuracy but lack sufficient biological insight [9], our proposed regularized multinomial logistic regression framework yields models that are both transparent and robust. These advantages are especially important in small-sample clinical studies where complex models are prone to overfitting due to insufficient training data and interpretable gene signatures are critical for clinical relevance [10].

Across multiple external datasets, our SGL framework consistently achieved classification accuracy comparable to, and sometimes exceeding, that of other interpretable prediction methods such as multinomial logistic regression with lasso penalty [1] and random forest models [11]. Beyond predictive performance, the resulting models highlighted coherent sets of genes within disease-relevant cell types. These patterns closely align with previously reported biological findings in peer-reviewed studies, thereby providing direct insight into the cellular architecture underlying disease. By simultaneously delivering robust prediction and interpretable, cell-type-specific gene signatures that inform disease mechanisms, our SGL framework bridges a key gap in current single-cell disease classification approaches.

## 2 Methods

Our workflow consists of four major steps which we will discuss in detail in what follows: (i) construction of patient-level gene-by–cell-type pseudobulk RNA expression matrices, (ii) library-size normalization and highly variable gene selection, (iii) hyperparameter tuning for the SGL model using cross validation on the training data, and (iv) fitting the final SGL model with the selected hyperparameters and evaluating predictive performance on held-out samples. All analyses were conducted in R version 4.5.3 [12], implementation of the SGL model relied on the the `msgl` library version 2.3.9 [13] for SGL model fitting, and random forest models were fitted using the caret version 7.0.1 package [14].

### 2.1 Gene-by-cell-type Pseudobulk Matrices Construction

For a dataset with *n* samples, *p* genes, and *L* cell types, we aggregate the raw single-cell RNA counts into patient-level pseudobulk profiles [5]. Specifically, for sample *i* and cell type *l*, let *C*_*igl*_ denote the raw count of the abundance of the gene *g* aggregated across all single cells in cell type *l*. For each sample *i*, we sum the *C*_*igl*_ across all genes and cell types, producing *p* × *L* features per sample. This results in a *n* × *pL* feature matrix where each block of *p* columns corresponds to one cell type. This representation preserves cell-type specificity while enabling direct use of classical supervised learning methods.

### 2.2 Library-size Normalization

To correct for differences in sequencing depth across samples before downstream analyses, we applied three commonly used library-size normalization strategies: Counts-Per-Million (CPM) [15], Upper Quartile (UQ) [16], and Median Ratio (MR) [17] (see Supplementary Text for details). In all cases, normalization was performed by computing cell-type-specific size factors, allowing each cell type to be scaled independently and thereby accounting for heterogeneity in expression distributions across cell types.

These normalizations could alternatively be applied to the full pseudobulk matrix, across all *p* × *L* features without distinguishing cell types. However, by normalizing separately within each predefined cell type block, we remove effects due to differential cell type composition. While such differences could be predictive, we made this choice so that the significance of genes and cell types would be interpretable as due to gene expression differences.

Following normalization, counts were then log_2_-transformed with a pseudo-count of 1, i.e., each normalized count *x* was replaced by log_2_(*x*+1), to reduce skewness, stabilize variance, and mitigate the dominance of highly expressed genes in downstream analyses.

### 2.3 Highly Variable Gene Selection

After library-size normalization, the top *N* highly variable genes (HVGs) [18] are selected based on the log-normalized pseudobulk data to identify the most informative features for downstream modeling. We implemented in our package scSGL two alternative strategies depending on user preference: a DESeq2-based [19] variance-stabilizing transformation approach (`hvg_deseq`) and a variance-based naive selection approach (`hvg_naive`). For `hvg_deseq`, we follow the standard DESeq2 workflow: low-expression genes with counts below a specified threshold (usually set to 5-10, the default being 5) in more than *N* − 10 samples are removed, retaining only those expressed above the threshold in at least 10 samples. Variance-stabilizing transformation is then applied, and the top *N* features by variance are retained. For `hvg_naive`, features are ranked by variance across samples. The top *N* genes are selected.

Importantly, HVG selection can be performed either jointly across all features (whole-space) or separately within each cell type (`per-cell-type`) to capture cell type–specific variability. However, applying across all features could result in input features a priori dominated by certain cell types, which can make interpretability of the importance of cell types problematic.

For the results reported in the paper, we used `per-cell-type` HVG selection to avoid being dominated by cell types with more abundant expression counts, and chose the `hvg_deseq` strategy to produce a stable variance estimate. In total, we picked the top *N* = 3000 HVGs per cell type based on the normalized pseudobulk matrices.

### 2.4 Multinomial Logistic Regression with Sparse Group Lasso Penalty

Consider a multiclass classification problem with *K* outcome classes, *n* samples, *L* cell types. For each sample *i* = 1, …, *n*, let 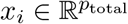 denote its pseudobulk gene-by-cell-type expression profile, and let *y*_*i*_ ∈ *{*1, …, *K}* be the class label.

We follow the symmetric multinomial parametrization of the `msgl` package [13], in which all *K* classes are treated equally. Let 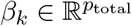 and 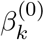 denote the coefficient vector and intercept for class *k*. The class probabilities are

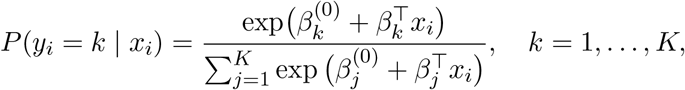

and the negative log-likelihood can then be computed as

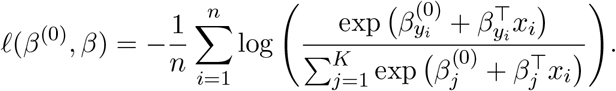

While this symmetric formulation is overparameterized under an unpenalized setting, the sparse group lasso penalty yields a well-defined solution by shrinking coefficients toward zero, making an explicit reference-class constraint unnecessary. This formulation also avoids imposing an artificial advantage or disadvantage on any particular class, ensuring that the penalty operates uniformly across all *K* classes.

The sparse group lasso penalty [8] imposes a group structure by grouping the coefficients according to *a priori* groupings of the feature. We use this framework where the grouping of our features corresponds to cell types. For each cell type *l* = 1, …, *L*, let *p*_*l*_ denote the number of genes included for that cell type after preprocessing. The total number of coefficients is 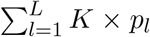, since each gene contributes a coefficient for each of the *K* classes. We collect the coefficients for cell type *l* into a submatrix 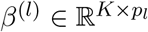, where row *k* corresponds to class *k* ∈ *{*1, …, *K}* and column *g* corresponds to gene *g* ∈ {1, …, *p*_*l*_}. We denote by 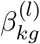 the coefficient of gene *g* in cell type *l* for predicting class *k*.

The sparse-group lasso penalized objective of [8] adds the following penalty to *ℓ*(*β*^(0)^, *β*),

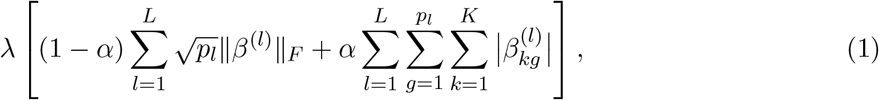

where 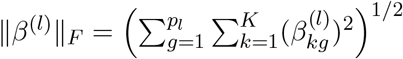.

Here *λ >* 0 controls the overall regularization strength, and *α* ∈ [0, 1] trades off cell-type–level sparsity (group lasso penalty) and gene-level sparsity (lasso penalty). The 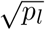 factor balances penalization across cell types with differing numbers of features / genes. When *α* = 1, the penalty reduces to a standard lasso [1], which we include as a baseline comparison.

### 2.5 Data Summary

To evaluate the predictive performance and interpretability of our SGL framework against standard lasso and random forests, we assessed its application to three diverse single-cell datasets spanning autoimmune disease, viral infection, and cancer. We selected these datasets to represent distinct biological contexts for classification: systemic immune dysregulation (where immune cells are the main drivers) and solid tissue pathology (where tumor cells drive the disease signals).

The systemic lupus erythematosus SLE [20] dataset profiles over 1.2 million peripheral blood mononuclear cells (PBMCs) from 205 patients and 131 controls. The COVID-19 [21] dataset profiled COVID, non-COVID, healthy control, and lipopolysaccharide (LPS)-simulated healthy PBMC samples across three research sites at Cambridge, Newcastle, and the Sanger Institute. The colorectal cancer CRC [22] dataset includes 99 tumor and adjacent normal samples with patients’ tumors classified as belonging to either mismatch repair-deficient (MMRd) or mismatch repair-proficient (MMRp) subtypes. Together, these datasets provide diverse biological settings where disease states are driven by coordinated cell-type-specific gene expression patterns, creating a platform on which we can evaluate whether SGL can recover biologically meaningful signatures while maintaining strong predictive performance. See Supplementary Text for more information on these datasets.

Table 1 summarizes the sample distribution for all three datasets. As shown, these real-world clinical datasets exhibit different degrees of class imbalance and batch-specific heterogeneity. For instance, in the SLE dataset, phenotypes such as Flare and Treated are completely absent from batches 1, 2, and 4. Similarly, with the COVID-19 dataset, LPS and Non-COVID samples are missing from Cambridge and Sanger sites. To address this imbalance and ensure a more robust evaluation, we restrict our analysis to the dominant conditions within each dataset. For SLE dataset, we focused on classifying two major subtypes Managed and Normal; for COVID-19 dataset, we focused on distinguishing COVID patients and healthy controls; for CRC dataset, we performed binary classification of tumor subtypes MMRd and MMRp.

**Table 1:**
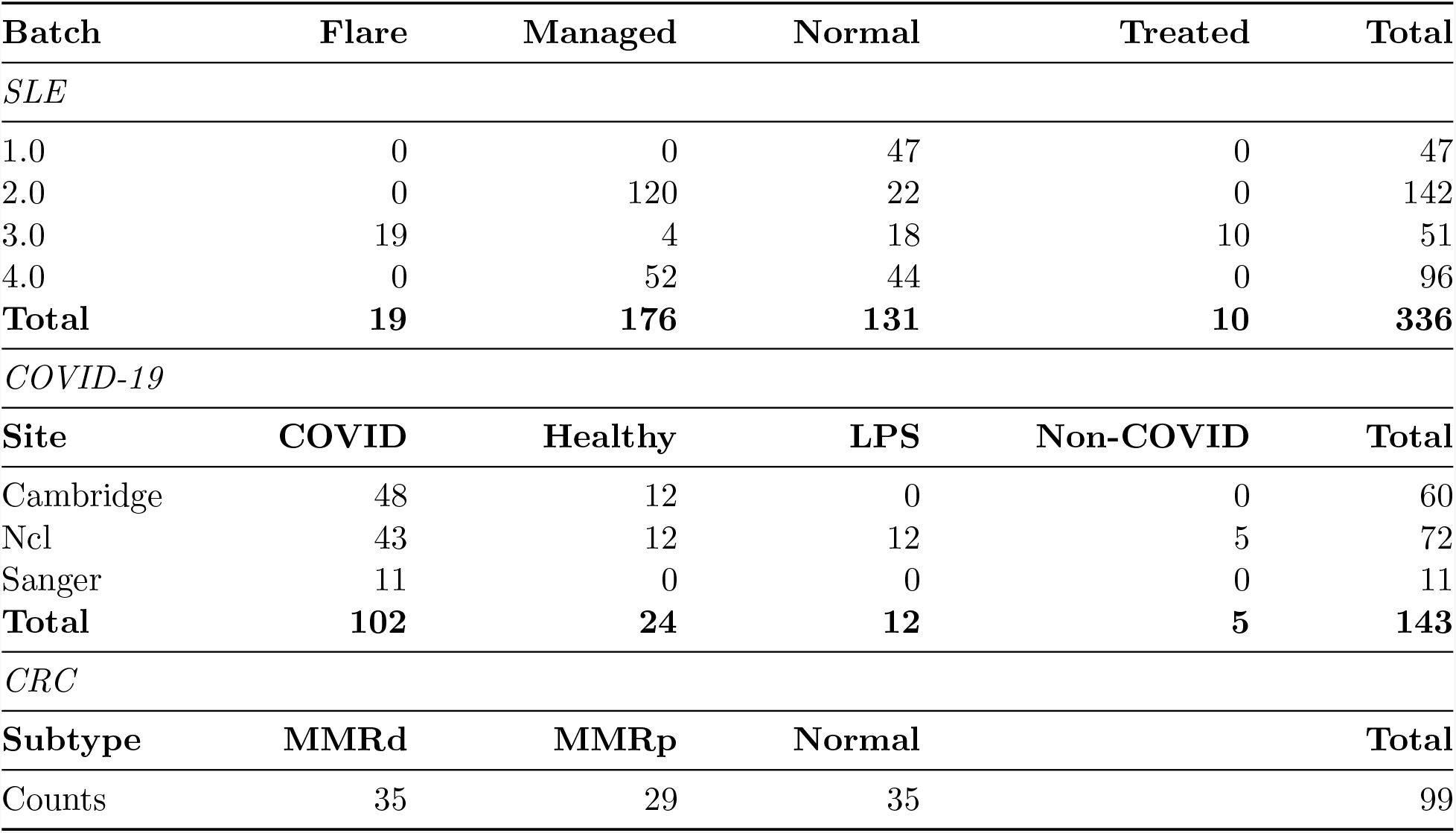
Condition distribution across the three datasets (SLE, COVID-19, CRC).

### 2.6 Train-Test Splitting and Cross-Validation

To evaluate predictive performance, we partitioned each dataset into training and independent test sets using an 80–20 split at the sample level, stratified by disease condition to preserve class proportions. For the SLE dataset, we additionally examined a second training strategy in which models were trained exclusively on Batch 2 — the largest batch containing both Managed and Normal samples — and evaluated on the remaining batches, in order to assess model robustness to batch heterogeneity (See Section 3.4).

To prevent information leakage during model evaluation, all preprocessing decisions—including normalization scaling factors, highly variable gene selection, and hyperparameter tuning—were estimated exclusively from the training data at each stage, both within cross-validation folds and before final evaluation on the held-out test set. The held-out test set was accessed only once after all model training was complete to estimate out-of-sample predictive performance.

For the SGL models, we performed nested 10-fold cross-validation. The outer loop evaluated all three normalization methods (UQ, CPM, MR), the middle loop varied the lasso to group lasso regularizing parameter

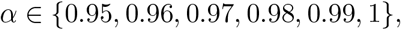

and the inner loop intended to optimize the regularization strength over a grid of 20 *λ* values spaced evenly on the log scale between 0.01 and 0.2 using the `msgl::fit` function. Note that *α* = 1 translates to standard lasso model [1] which will be used for baseline comparison.

Cross-validation selected the following optimal combinations:

- **SLE (Batch 2 only):** MR normalization, *α* = 0.95, *λ* = 0.018.
- **SLE (80% random sampling):** UQ normalization, *α* = 0.96, *λ* = 0.012.
- **CRC:** UQ normalization, *α* = 0.98, *λ* = 0.012.
- **COVID-19:** UQ normalization, *α* = 0.98, *λ* = 0.090.

We also performed nested 10-fold cross-validation to identify the optimal random forest models [11] for comparisons. The outer loop evaluated the same three normalization methods (UQ, CPM, MR), while the inner loop used `caret::train` function to tune the number of predictors sampled at each split, evaluating

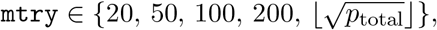

*p*_total_ denotes the total number of pseudobulk features after HVG selection step. As with the SGL models, both training strategies were examined for the SLE dataset (Batch 2 only and 80% random sampling), whereas for the COVID-19 and CRC datasets cross-validation was conducted on an 80% random split of the entire dataset.

Cross-validation selected the following optimal normalization methods for the random forest models:

- **SLE (Batch 2 only):** MR normalization.
- **SLE (80% random sampling):** CPM normalization.
- **COVID-19:** MR normalization.
- **CRC:** CPM normalization.

### 2.7 Feature Selection and Importance Score Thresholding

To identify the most predictive features from each model for downstream interpretability analysis, we applied method-specific thresholds to the fitted model outputs. For SGL and lasso models, we retained features whose absolute standardized coefficient values exceeded 0.02 at the CV-selected *λ*. For random forest models, importance scores measure each feature’s average contribution to reducing classification error across decision trees [11]. We retained features whose importance scores exceeded a threshold of 60. For pseudobulk differential gene expression analysis using `DESeq2` [19], we retained features with adjusted *p*-values below 0.05.

## 3 Results

### 3.1 Summary of the SGL Approach

We model patient disease under a high-dimensional, sparse multinomial logistic regression framework that utilizes gene-by–cell-type pseudobulk features, where each sample is represented by a concatenated vector of gene expression summaries across all predefined cell types. This formulation allows the model to simultaneously incorporate gene level and cell-type–specific information when making predictions.

To leverage this biological structure, we apply the sparse group lasso (SGL) penalty [8], with groups defined as the group of gene expressions coming from a particular cell type. Unlike the standard lasso [1] that penalizes all features equally, SGL induces a structured regularization that operates on two levels: the penalty encourages the model to select relevant cell types while setting coefficients for non-informative cell types to zero; within selected cell types, the penalty further imposes sparsity at the individual gene level. This structured selection yields coherent cellular signatures that form the basis for both the strong predictive performance and the high interpretability observed in subsequent analyses.

### 3.2 Prediction accuracy

We evaluated the predictive performance of our proposed SGL framework across the SLE, COVID-19, and CRC datasets, comparing it against two baselines which also provide interpretability of input features: multinomial logistic regression with a standard lasso penalty [1] and random forest (RF) [11] classifiers. Models were developed using an 80–20 train-test split. We used nested 10-fold cross-validation on the training set to select optimal hyperparameters, after which the final model was retrained on the complete training set and assessed on the held-out test set (see Section 2.6 for details).

Table 2 summarizes the testing accuracy under an 80/20 train–test split for the Sparse Group Lasso (SGL), standard lasso, and random forest (RF) across the three datasets.

**Table 2:** Testing accuracy of the Sparse Group Lasso (SGL) framework compared to standard lasso and random forest (RF) under an 80/20 train-test split for the SLE, CRC, and COVID-19 datasets.

| Data Set | Sample Number | SGL | Lasso | RF |
| --- | --- | --- | --- | --- |
| *SLE* |  |  |  |  |
| Normal | 26 | 0.73 | 0.85 | 0.85 |
| Managed | 35 | 0.97 | 0.97 | 0.94 |
| *CRC* |  |  |  |  |
| MMRd | 7 | 0.86 | 0.71 | 0.86 |
| MMRp | 5 | 0.80 | 0.80 | 0.60 |
| *COVID-19* |  |  |  |  |
| COVID | 20 | 0.95 | 0.95 | 0.95 |
| Healthy | 4 | 0.50 | 0.25 | 0.25 |

For the SLE dataset, SGL achieves near-identical performance to lasso and RF in the Managed class (0.97 vs 0.97 and 0.94, respectively). Although SGL performs slightly lower in the Normal class (0.73 vs 0.85), its performance remains competitive, particularly considering the relatively small sample size (n = 26). Importantly, SGL maintains strong accuracy in the clinically relevant managed subgroup without sacrificing generalization. For the CRC dataset, in the MMRd subtype, SGL achieves 0.86 accuracy, outperforming lasso (0.71) and matching RF (0.86). In the MMRp subtype, SGL matches lasso (0.80) and substantially exceeds RF (0.60). For the COVID-19 dataset, SGL matches both lasso and RF in the COVID class (0.95 across all methods). Notably, in the Healthy class, SGL achieves 0.50 accuracy, doubling the performance of lasso and RF (0.25), though the sample size is limited (n = 4).

Collectively, these results demonstrate that SGL is a competitive and robust predictive model. It does not compromise the predictive accuracy of traditional approaches. In most settings, SGL either matches or exceeds the performance of lasso and RF while offering improved interpretability through its sparsity in both gene and cell type levels.

### 3.3 Interpretable features

The fitted coefficient values from the our SGL models provide a direct and interpretable summary of disease-associated effects. Nonzero coefficients identify which cell types contribute to disease classification and, within those cell types, which genes drive the predictive signals. Larger absolute coefficient values for one disease subtype correspond to stronger predictive contributions to that class. Moreover, to further assess the stability of these signatures, we examined the coefficient paths of significant gene-by-cell-type signatures, defined as the trajectory of coefficient values as the regularization strength *λ* is relaxed for a fixed *α* value, indicating coordinated entry of gene features within these biologically meaningful cell types. We note that the coordinated entry and persistence of coefficients across a wide range of *λ* values is a consequence of the group penalty structure of SGL rather than independent evidence of biological relevance. Nonetheless, examining coefficient paths provides a useful diagnostic for understanding which cell types and genes the model prioritizes, and the biological relevance of the selected features can be assessed by comparison with known disease biology as we will demonstrate next.

Building on this interpretation, we first compared coefficient paths obtained from SGL model (*α* = 0.96) to those from the standard lasso model (*α* = 1) in predicting Managed class on the SLE dataset in Figure 1. Under the SGL penalty, trajectories of coefficients exhibit a structured and biologically coherent pattern that nonzero coefficients emerge predominantly within three immune cell types classical monocytes, CD4^+^ T cells, and CD8^+^ T cells. These findings align well with known SLE biology as they have been established as main drivers of SLE pathogenesis [20, 23, 24]. The SGL penalty optimizes a joint loss function over group and individual levels and induces sparsity at both levels simultaneously. Consequently, as the regularization strength decreases, genes within these selected cell types enter the SGL model in a coordinated manner, reflecting the group-level structure imposed by the SGL penalty. This behavior reflects the objective of SGL penalty, to identify a predictive set of cell types and their specific gene drivers through joint regularization, rather than treating genes as isolated features.

**Figure 1:**
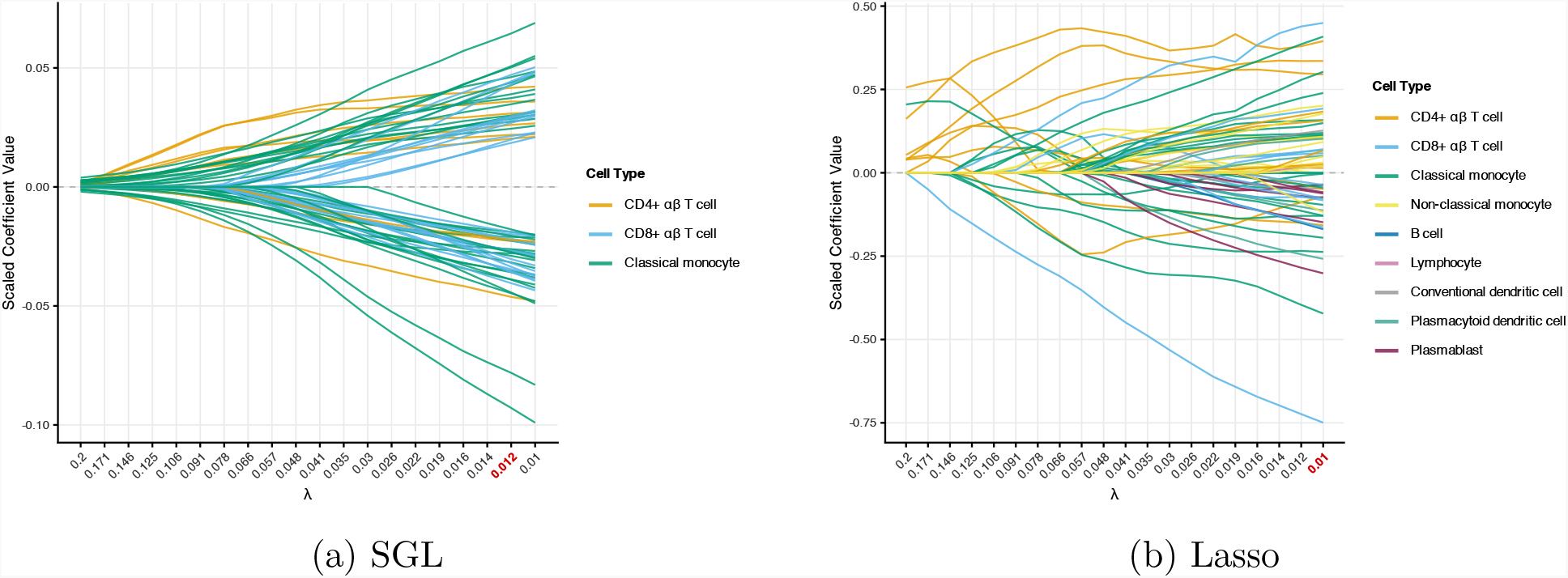
Coefficient paths for the SLE dataset (Managed condition) comparing Sparse Group Lasso (left) and standard Lasso (right). Features were filtered using the thresholds described in Section 2.7: only features with |coef| ≥ 0.02 at any *λ* are shown. The red tick mark indicates the CV-selected *λ* value.

On the other hand, coefficient paths produced by the standard lasso model are significantly less structured on the SLE dataset. Nonzero coefficients appear scattered across a wide range of cell types, with no clear concentration on specific immune populations. This fragmented selection pattern reflects the well-known behavior of the lasso in the presence of correlated predictors: when multiple genes within a cell type are correlated, the lasso tends to select one arbitrarily among them [7], capturing incoherent gene signatures that obscure the underlying cellular organization of disease. Therefore, while standard lasso models can achieve comparable predictive accuracy, its coefficients offer limited biological insight into which cell types drive disease classification, and often resulting in incredibly sparse gene sets that lack the coherence required for biological interpretation.

We can more carefully quantify these observations regarding the coefficient paths by focusing on the features selected by each final model according to the thresholding criteria described in Section 2.7. Figure 2 summarizes the cell type composition of the genes identified by four approaches: SGL, Lasso, Random Forests, and differential expression analysis. The SGL-selected gene features are, as previously mentioned, strongly concentrated in classical monocytes, CD4^+^ T cells, and CD8^+^ T cells, which are well-established drivers of systemic lupus erythematosus pathogenesis [20, 23, 24]. While random forest importance scores also include a large portion of classical monocytes and CD4^+^ T cells immune populations, the selected features are spread across a much broader range of cell types including non-classical monocytes, natural killer cells, B cells, and conventional dendritic cells. This diffuse selection pattern reflects the fact that random forest evaluates feature importance independently without leveraging the group structure, and therefore lacks the focused cellular signatures that facilitate biological interpretation. Similarly, standard lasso also identifies genes from classical monocytes, CD4^+^ T cells, and CD8^+^ T cells as important drivers, but also selects genes that are diffusely distributed across multiple immune cell types, reflecting its tendency to select correlated predictors in a unstructured manner. Consistent results are observed when no coefficient filtering is applied (See Supplementary Figure S-9), where SGL continues to concentrate selected features in classical monocytes, CD4^+^ T cells, and CD8^+^ T cells, while lasso and random forest exhibit broader and less focused cell-type selection patterns.

**Figure 2:**
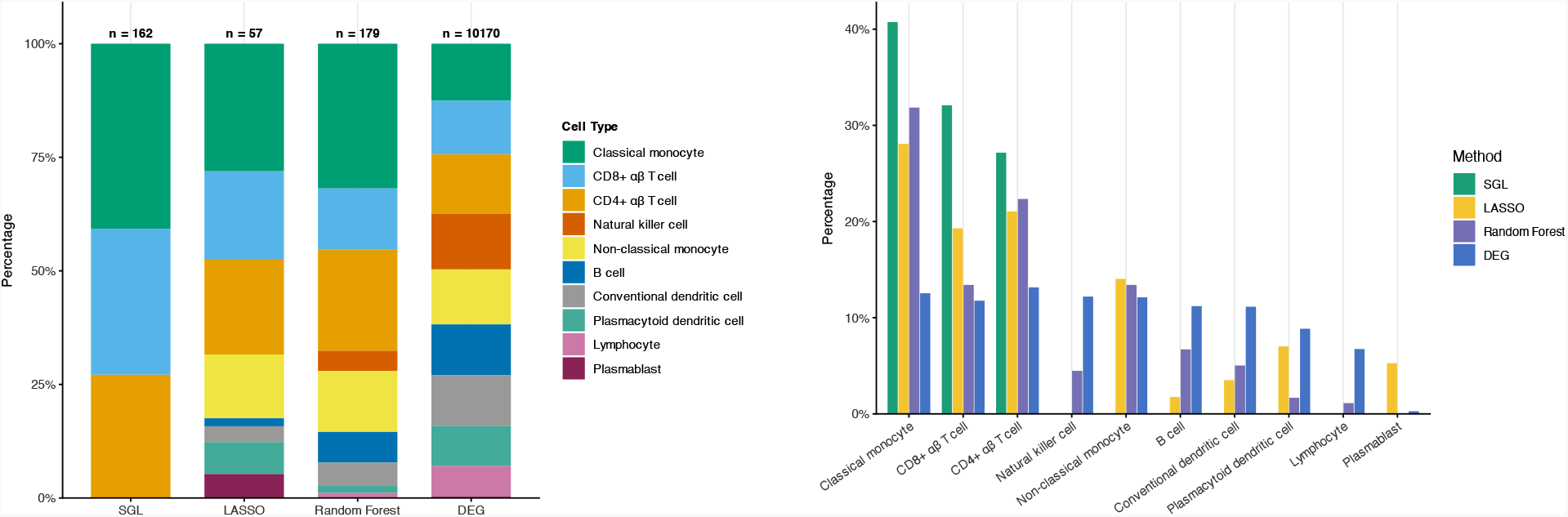
Cell type composition plots of important features for the SLE dataset comparing SGL, Lasso, Random Forest, and Differential Gene Analysis. Features were filtered using the thresholds described in Section 2.7: |coef| ≥ 0.02 for SGL and lasso, and importance score ≥ 60 for random forest.

Similar patterns are observed in the CRC and COVID-19 datasets (Supplementary Figures S-10, S-11, S-12, and S-13). For the COVID-19 dataset, SGL concentrates its selected features in CD14^+^ monocytes, plasmablasts, as well as CD16^+^ monocytes, consistent with the central role of CD14^+^ and CD16^+^ monocytes in driving the COVID-19 immune response [21], while lasso and random forest select features distributed across a much broader range of cell types with no clear concentration in disease-relevant populations. For the CRC dataset, all methods concentrate on epithelial cells, which is expected as MMRd and MMRp subtypes are primarily distinguished by tumor epithelial gene expression differences [22].

Figure 3 shows the overlap of selected features among SGL, lasso, and random forest on the SLE dataset. The three methods identify largely distinct predictive feature sets, with only 28 features shared across all methods. Together with the previous Figure 2, this suggests that the structured regularization of SGL identifies a different and more biologically coherent set of disease-associated signals compared to lasso and random forest. Similar patterns were observed in the COVID-19 and CRC datasets (Supplementary Figures S-14, S-15).

**Figure 3:**
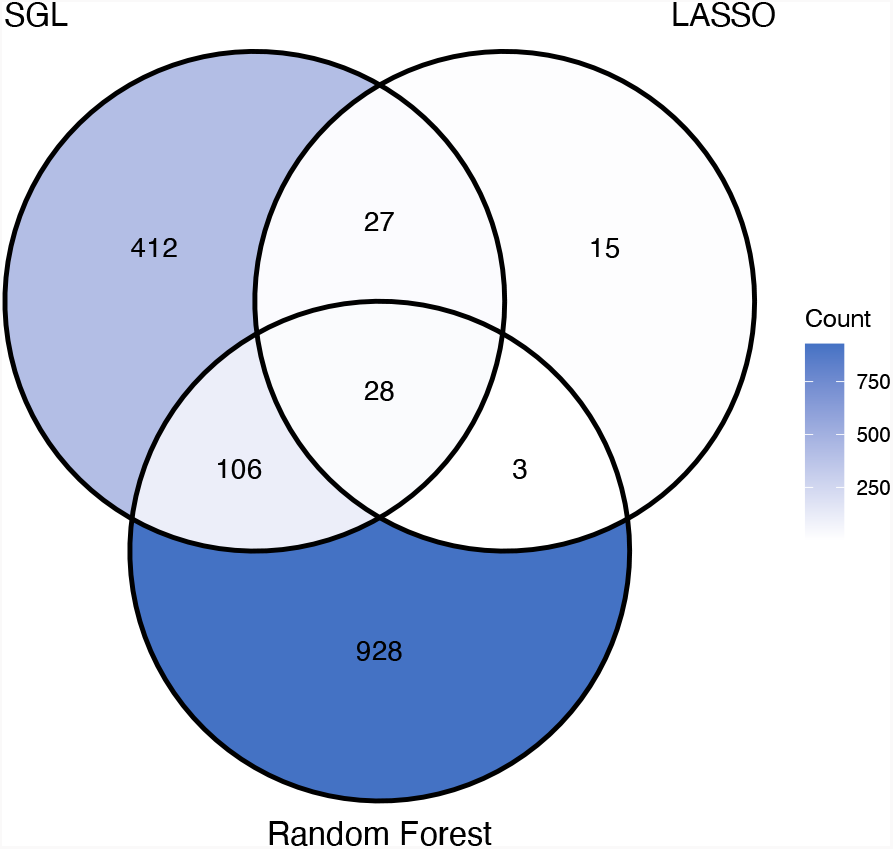
Feature overlap across three methods (SGL, Lasso, Random Forest) for the SLE dataset. Features were filtered using the thresholds described in Section 2.7: |coef| ≥ 0.02 for SGL and lasso, and importance score ≥ 60 for random forest.

We can also compare to other strategies that can be used to identify relevant cell-types. Pseudobulk differential expression (DE) analysis tests for differences between pseudobulk measures of gene expression within cell types; the number and strength of gene expression differences is often used informally to identify important cell-types [5]. Using `DESeq2` [19], DE analysis between `Managed` and `Normal` conditions in the SLE dataset identifies large numbers of significant genes broadly across nearly all cell types (Figure 2, Supplementary Table 4), intrinsically making it difficult to isolate which cell populations are most predictive of disease status. We observed a similar lack of prioritization when running DE between the `COVID` and `Healthy` conditions in the COVID-19 dataset (Supplementary Figures S-13, Supplementary Table 5). For the CRC dataset, DE between `MMRd` and `MMRp` conditions put prioritization on three cell types, including the predictive epithelial population (Supplementary Figures S-11, Supplementary Table 6).

Another strategy, Augur [25], is a cell type prioritization method for single-cell data that ranks cell types according to how accurately experimental conditions (e.g., disease states) can be predicted from gene expression profiles within each cell type. For each cell type, Augur trains a random forest classifier on repeated subsamples of single cells and uses cross validation performance quantified by AUC values as a measure of condition-associated transcriptional change. Cell types with higher predictive accuracy (higher AUC values) are interpreted as being more transcriptionally perturbed with respect to the condition of interest, reflecting larger and more consistent gene expression shifts. Notably, unlike all of the other strategies, Augur evaluates disease-associated perturbation at the single-cell level within each cell type independently and does not incorporate patient-level aggregation. For the SLE dataset, Augur failed to distinguish any single immune cell type with strong predictive power, with all cell types’ AUC values ranging narrowly between 0.503 and 0.582 (Supplementary Table 7). The highest ranked cell types, such as B cells and CD8-positive, alpha-beta T cells, exhibit only marginally higher AUC values than the remaining cell types. This narrow spread of AUC values indicates limited separation between responsive and non-responsive populations under Augur’s framework for the SLE dataset. Similarly, for the CRC dataset (Supplementary Table 9), the overall AUC values are low and tightly clustered between 0.515 and 0.589, with epithelial cells ranked highest but with a low AUC value 0.589, suggesting limited transcriptional separation at the single-cell level between MMRd and MMRp subtypes under Augur. In contrast, for the COVID-19 dataset (Supplementary Table 8), Augur successfully identified CD16^+^ monocytes, plasmablasts, and CD14^+^ monocytes as the three most highly perturbed cell types, consistent with our SGL discovery (Supplementary Figure S-13).

Taken together, these results demonstrate that the sparse group lasso framework provides a clear interpretability advantage. By leveraging the coordinated structure of our gene-by-cell-type features, SGL localizes disease-relevant signals in SLE dataset to specific immune populations while maintaining stable gene-level signatures within those cell types.

### 3.4 Handling Batch heterogeneity

While our previous evaluations utilized randomized 80/20 train-test splits, such standard practices can inadvertently mask the challenge of batch heterogeneity, a prevalent issue in single-cell genomics where technical variation confounds biological signals [26]. To rigorously assess the robustness of SGL against distribution shifts across batches, we conducted an experiment on the SLE dataset where we trained models exclusively on Batch 2 (which contains the majority of `Managed` and Normal samples) and evaluated their performance on the remaining batches (Table 3). The SLE dataset is well-suited to study batch heterogeneity because, for example, batch 3 has representation of each condition but with a limited of samples, and few Managed and Normal samples, while batch 1 only contains Normal samples.

**Table 3:** Testing accuracy of the Sparse Group Lasso (SGL) framework compared to standard lasso and random forest on the SLE dataset, when trained exclusively on Batch 2 (conditions managed and normal only).

| Test Set | Size | SGL | Lasso | RF |
| --- | --- | --- | --- | --- |
| Batch 1 - Normal | 47 | 0.81 | 0.87 | 0.64 |
| Batch 3 - Managed | 4 | 1.00 | 1.00 | 1.00 |
| Batch 3 - Normal | 18 | 0.39 | 0.50 | 0.39 |
| Batch 4 - Managed | 52 | 1.00 | 1.00 | 1.00 |
| Batch 4 - Normal | 44 | 0.14 | 0.09 | 0.25 |

The results of this experiment demonstrate that SGL maintains competitive robustness even when facing these challenging conditions (Table 3). For the `Managed` subtype, all three models (SGL, lasso, and random forest) achieved perfect accuracy (1.00) on Batches 3 and 4. Generalizing the `Normal` phenotype proved significantly more difficult due to technical drift. On Batch 1, SGL achieved an accuracy of 0.81 on normal samples, closely tracking lasso’s performance (0.87) and significantly outperforming the random forest (0.64). On Batch 3, performance dropped across all methods for normal samples, with SGL (0.39) performing comparably to random forest (0.39) but below lasso (0.50). On Batch 4, performance on normal samples declined further across all methods, reflecting a severe batch effect, with SGL (0.14) marginally outperforming lasso (0.09) but underperforming random forest (0.25). Taken together, neither SGL nor lasso uniformly performs better than the other across batches. This is consistent with the overall pattern observed in the 80–20 random split experiments, where no single method was uniformly superior across all conditions and datasets.

Furthermore, despite variability in predictive performance across batches, the interpretability of the SGL model remained stable. Features selected during this batch-restricted training continued to highlight the same key immune cell types — classical monocytes and T cells — as the primary drivers (Supplementary Figure S-16), consistent with established SLE biology [20, 23, 24]. In contrast, the standard lasso model exhibited a rather unstable feature selection across training strategies: when trained on Batch 2 only, lasso produced more near-zero scaled coefficients, whereas when trained on the 80% random split, it selected a more diffuse set of features spanning additional cell types including B cells, lymphocytes, conventional dendritic cells, and plasmablasts with larger coefficient magnitudes, which suggests that lasso’s feature selection is more sensitive to the composition of the training data, whereas SGL’s structured group penalty can help produce more consistent signatures. These results highlight that while batch effects and highly unbalanced class representation remain a significant hurdle for all single-cell classifiers, our SGL framework offers a robust alternative that perform comparatively with standard baselines in terms of prediction while preserving the biological coherence of its selected features.

## 4 Discussion / Conclusion

In this work, we introduced a Sparse Group Lasso (SGL) framework for the classification of disease states from single-cell RNA-seq data with multinomial logistic regression. Our primary contribution is demonstrating that predictive modeling in single-cell genomics need not to be be a trade-off between accuracy and interpretability. By imposing a structured penalty that respects both the cell type and gene hierarchies of the data, our model matches the predictive performance of classical methods (such as standard lasso and random forests) while yielding significantly more biologically coherent feature sets. As validated by differential expression analysis and established biology, the SGL framework successfully recovers known disease drivers, such as specific myeloid and lymphoid dysregulations in SLE and COVID-19 disease, offering a much more interpretable alternative to black box machine learning approaches.

Whole-sample single-cell data pose additional challenges related to extreme sparsity, technical artifacts, and computational scalability. Single-cell RNA-seq data typically exhibit an abundant amount of zero values due to dropout events and low capture efficiency, which exacerbates the risk of overfitting. Technical artifacts such as batch effects, platform heterogeneity, and variations in sample preparation pipelines all introduce noise that can confound true disease signals. Furthermore, processing datasets comprising hundreds of thousands of cells across patient cohorts demands algorithms that scale efficiently while remaining practical for routine clinical research. Although our framework does not explicitly address dropout events or correct batch effects at the cellular level, the pseudobulk strategy still substantially help reduce sparsity and enable stable, scalable inference.

Beyond its core performance, the regression-based nature of our framework offers considerable flexibility that was not fully explored in this study. Like any generalized linear model, the SGL formulation can be extended to control for clinical covariates such as demographic variables (e.g., age, sex) directly into the design matrix to adjust for potential confounders. Moreover, for interpretability-focused analyses, controlling for known technical factors may also help isolate biologically meaningful signals from the selected features. Additionally, while our current model relies on transcriptomic signatures, cellular abundance itself is often a hallmark of disease. The framework could be naturally augmented to include cell type composition (the number or proportion of cells per cell type) as additional predictors.

Taken together, we believe that this combination of competitive predictive performance and structured biological interpretability positions the SGL framework as a practical and principled tool for extracting biologically meaningful insights from single-cell data for patient-level classification.

