## Supplemental Figures and Tables for "A statistical framework for disease classification with scRNA-Seq Data"

#### Supplementary Figures and Tables

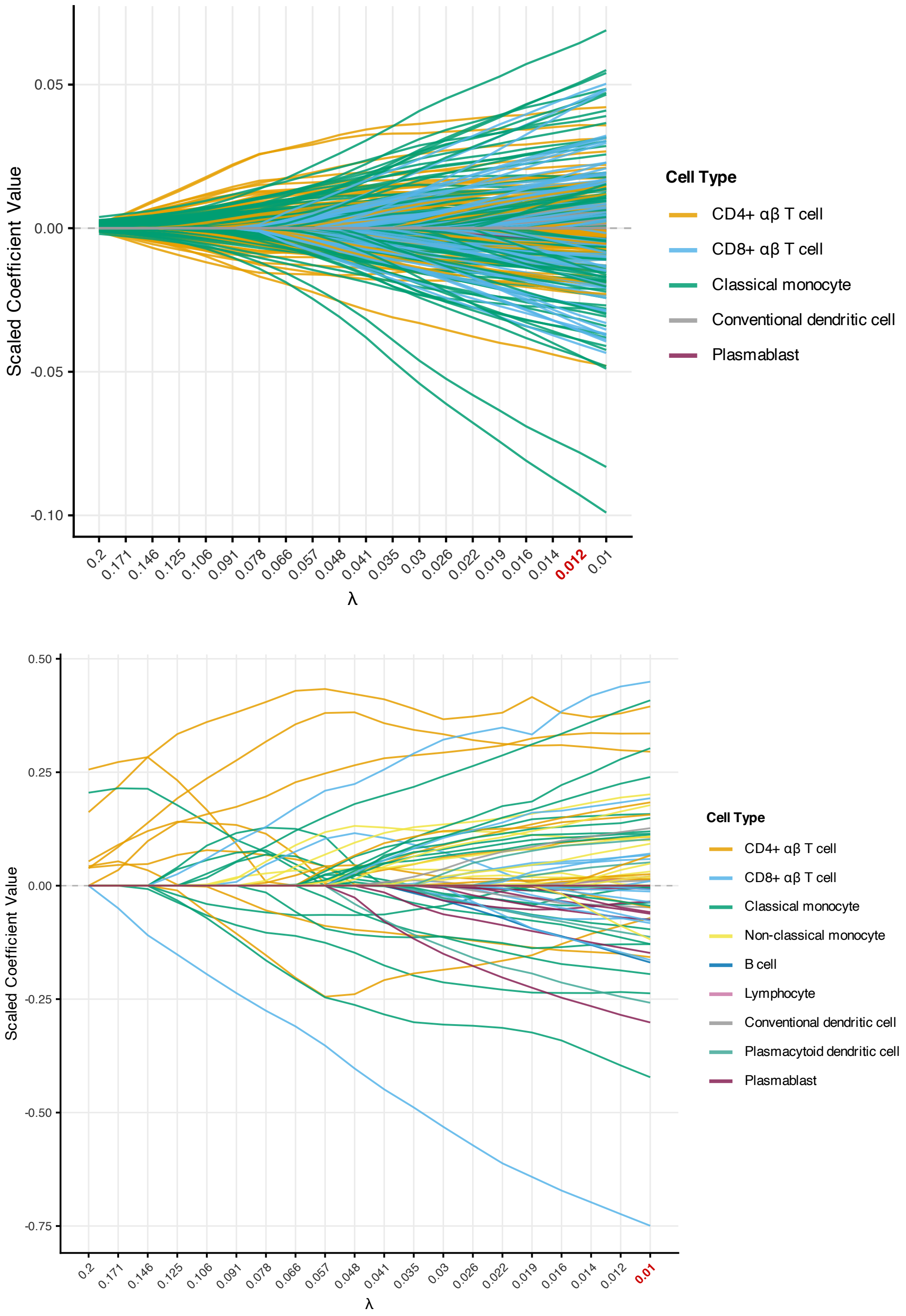


Figure S-1: Coefficient paths for the SLE dataset (Managed condition) comparing Sparse Group Lasso (top) and Lasso (bottom). Coefficients are not filtered.

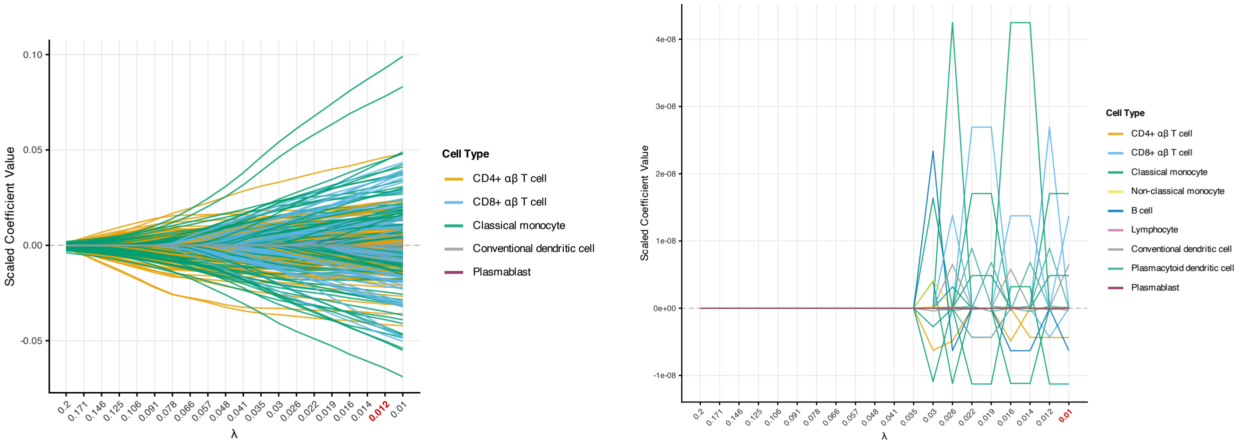


Figure S-2: Coefficient paths for the SLE dataset (Normal condition) comparing Sparse Group Lasso (left) and Lasso (right). Coefficients are not filtered based on their coefficient values.

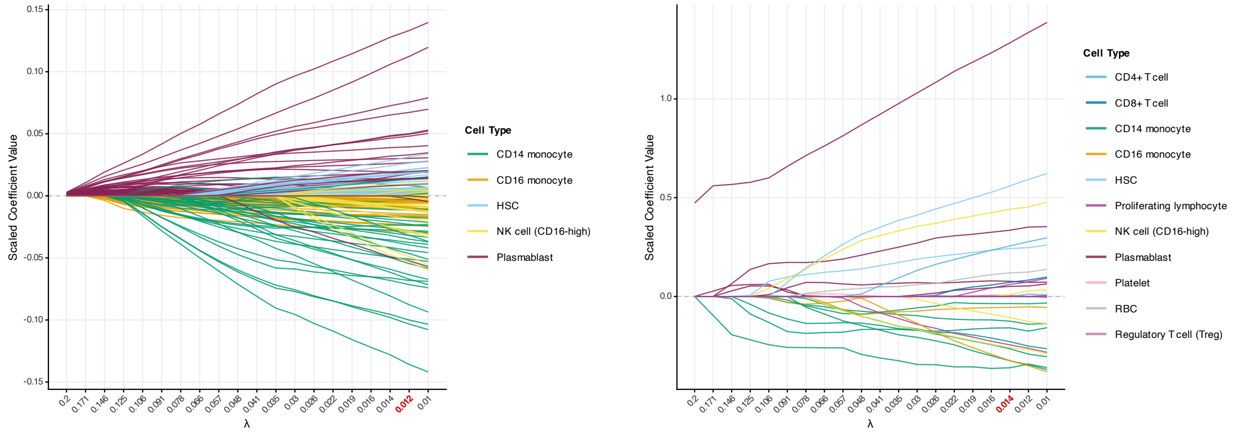


Figure S-3: Coefficient paths for the COVID dataset (Covid condition) comparing Sparse Group Lasso (left) and Lasso (right). Coefficients are not filtered.

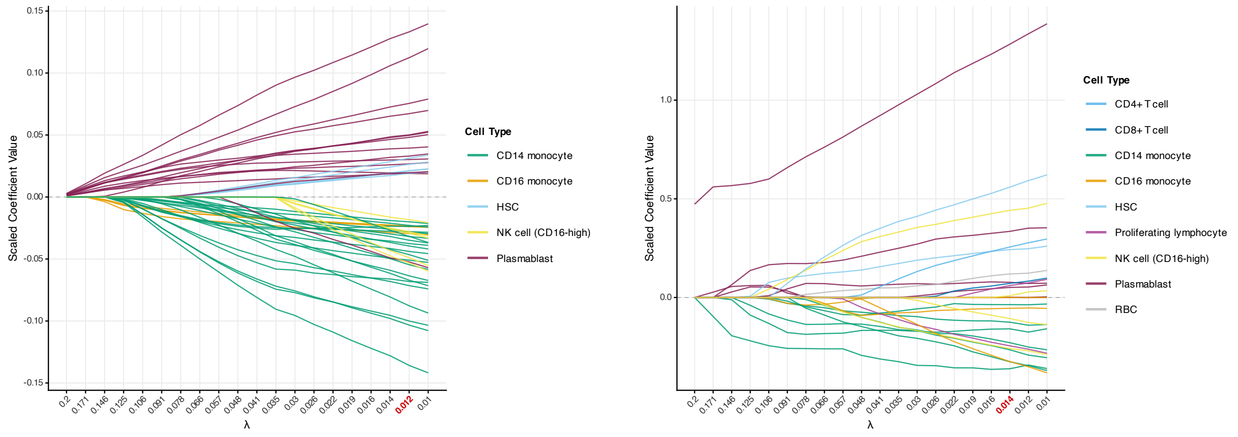


Figure S-4: Coefficient paths for the COVID dataset (Covid condition) comparing Sparse Group Lasso (left) and Lasso (right). Coefficients are filtered at 0.02.

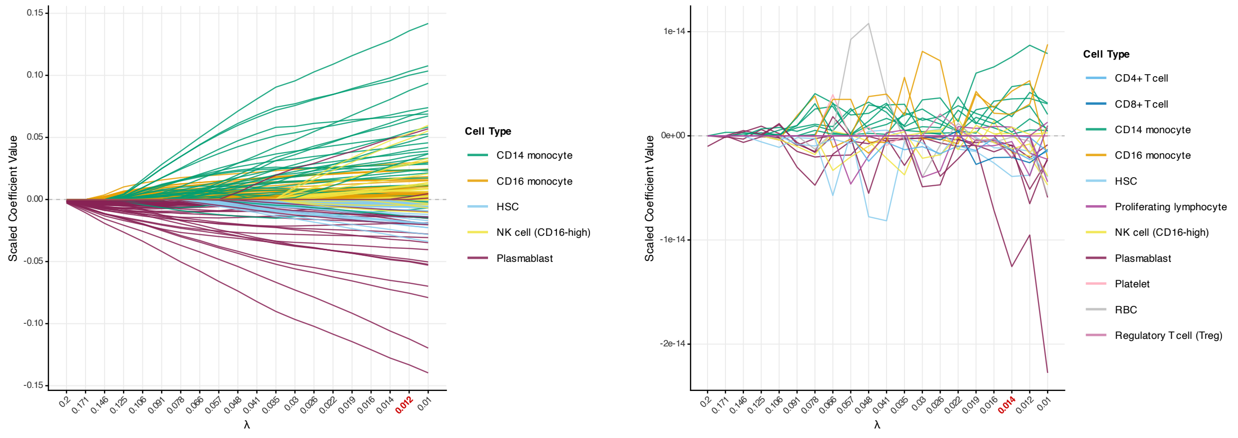


Figure S-5: Coefficient paths for the COVID dataset (Healthy condition) comparing Sparse Group Lasso (left) and Lasso (right). Coefficients are not filtered.

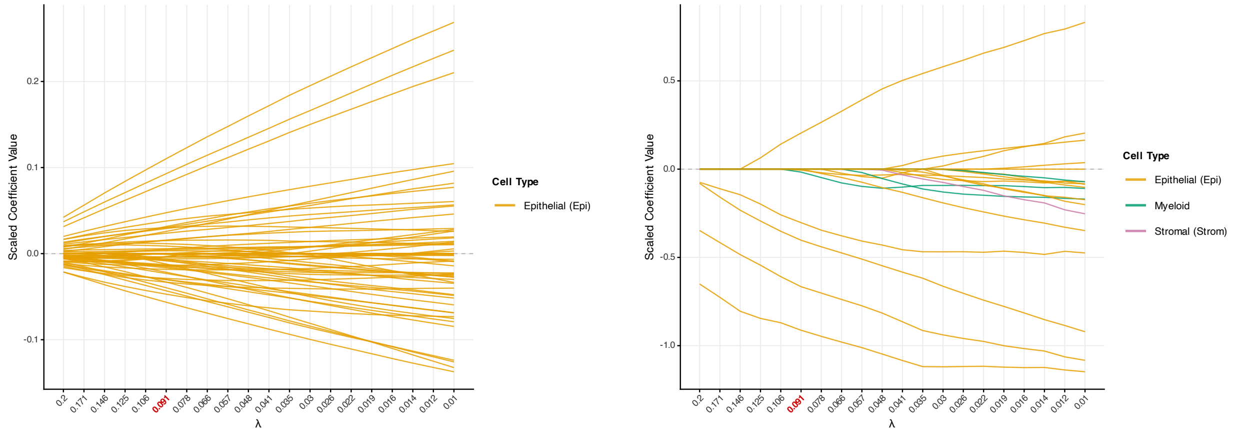


Figure S-6: Coefficient paths for the CRC dataset (MMRd condition) comparing Sparse Group Lasso (left) and Lasso (right). Coefficients are not filtered.

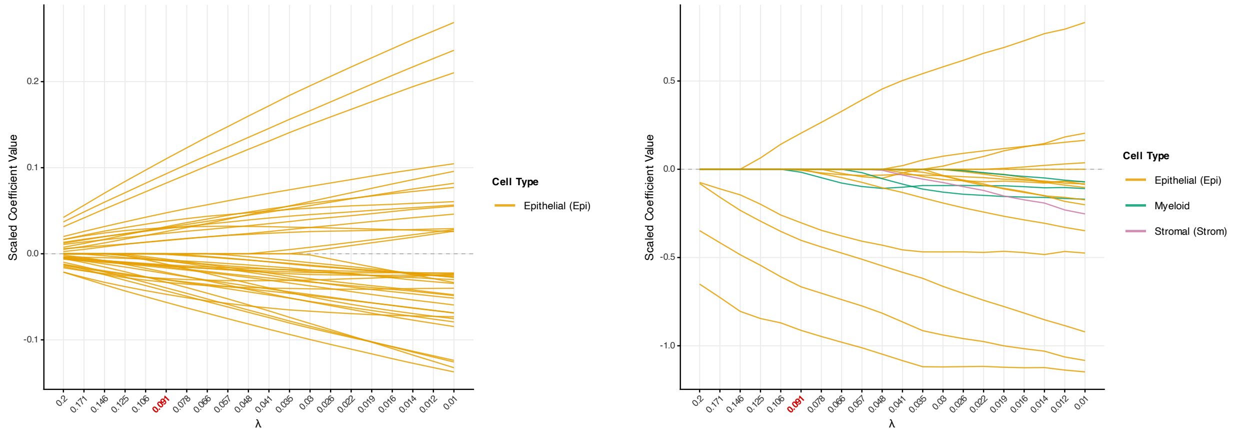


Figure S-7: Coefficient paths for the CRC dataset (MMRd condition) comparing Sparse Group Lasso (left) and Lasso (right). Coefficients are filtered at 0.02.

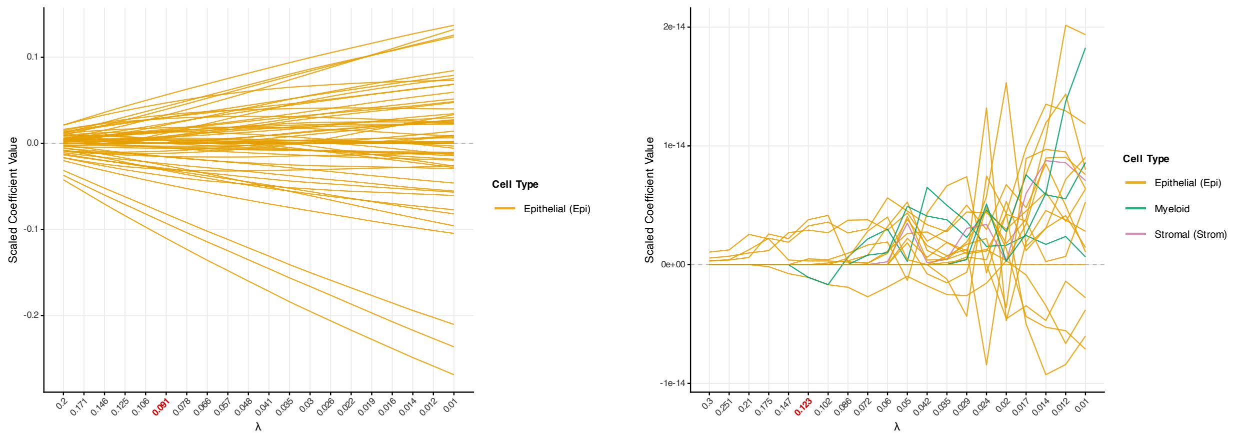


Figure S-8: Coefficient paths for the CRC dataset (MMRp condition) comparing Sparse Group Lasso (left) and Lasso (right). Coefficients are not filtered.

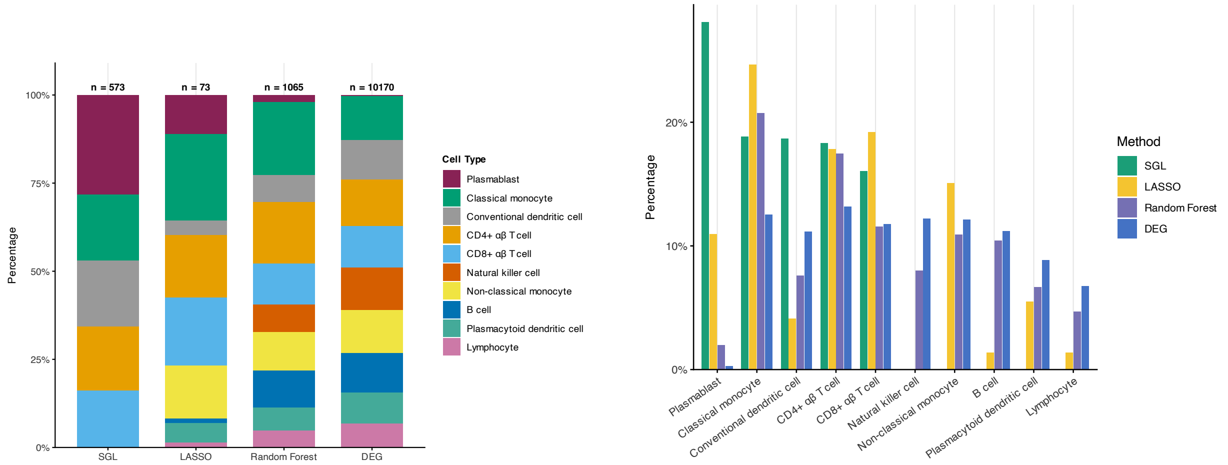


Figure S-9: Cell type composition plots of important features for the SLE dataset comparing SGL, Lasso, Random Forest, and Differential Gene Analysis. Random forest importance score filtered at 50 (baseline). SGL and Lasso coefficient values are not filtered. DEG adjusted p-values filtered at 0.05.

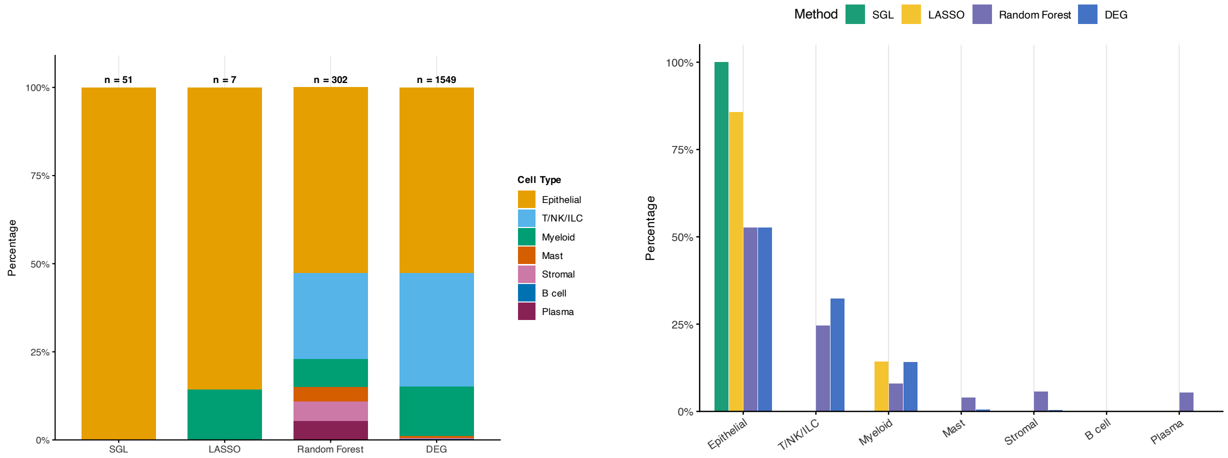


Figure S-10: Cell type composition plots of important features for the CRC dataset comparing SGL, Lasso, Random Forest, and Differential Gene Analysis. Random forest importance score filtered at 50 (baseline). SGL and Lasso coefficient values are not filtered. DEG adjusted p-values filtered at 0.05.

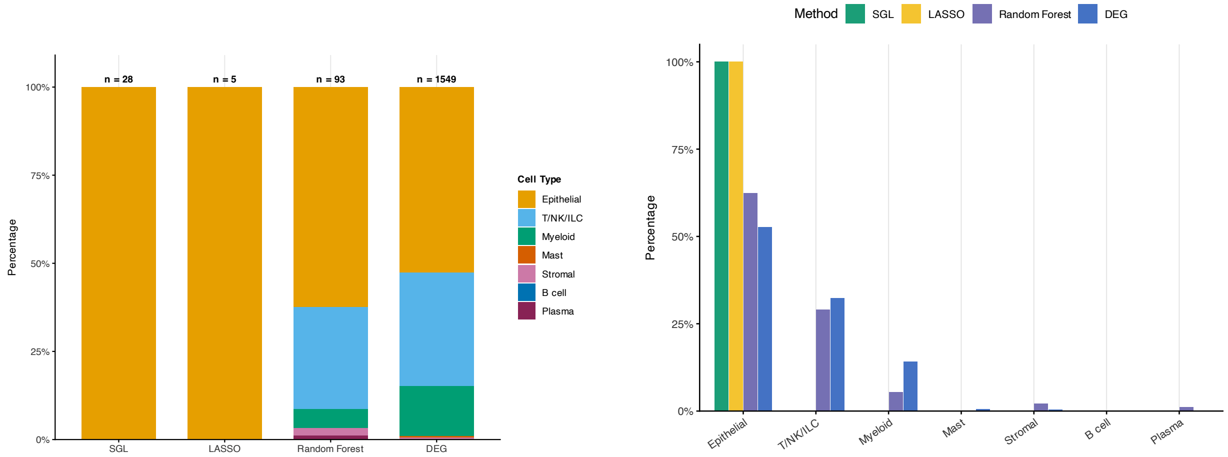


Figure S-11: Cell type composition plots of important features for the CRC dataset comparing SGL, Lasso, Random Forest, and Differential Gene Analysis. Random forest importance score filtered at 60. SGL and Lasso coefficient values filtered at 0.02. DEG adjusted p-values filtered at 0.05.

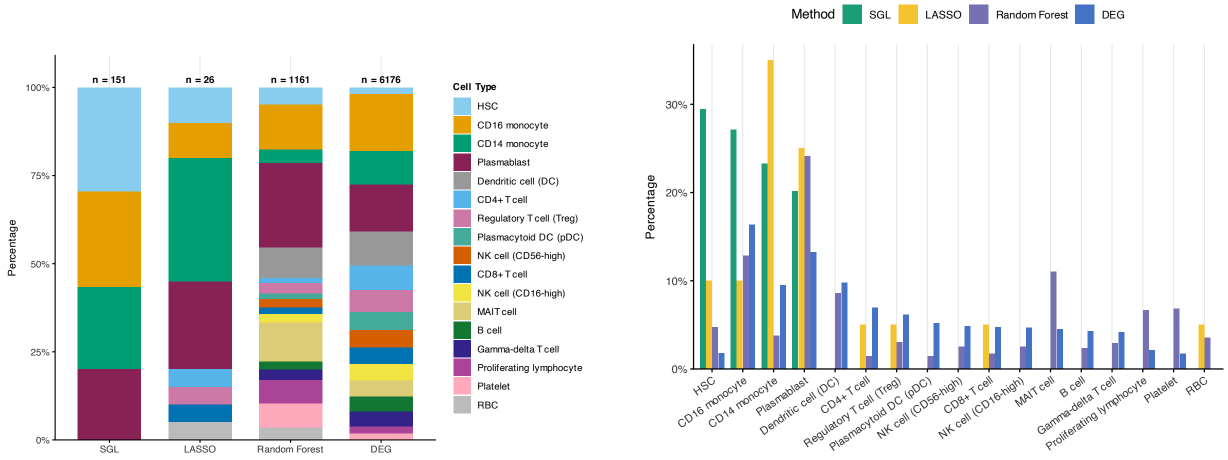

Figure S-12: Cell type composition plots of important features for the COVID dataset comparing SGL, Lasso, Random Forest, and Differential Gene Analysis. Random forest importance score filtered at 50 (baseline). SGL and Lasso coefficient values are not filtered. DEG adjusted p-values filtered at 0.05.

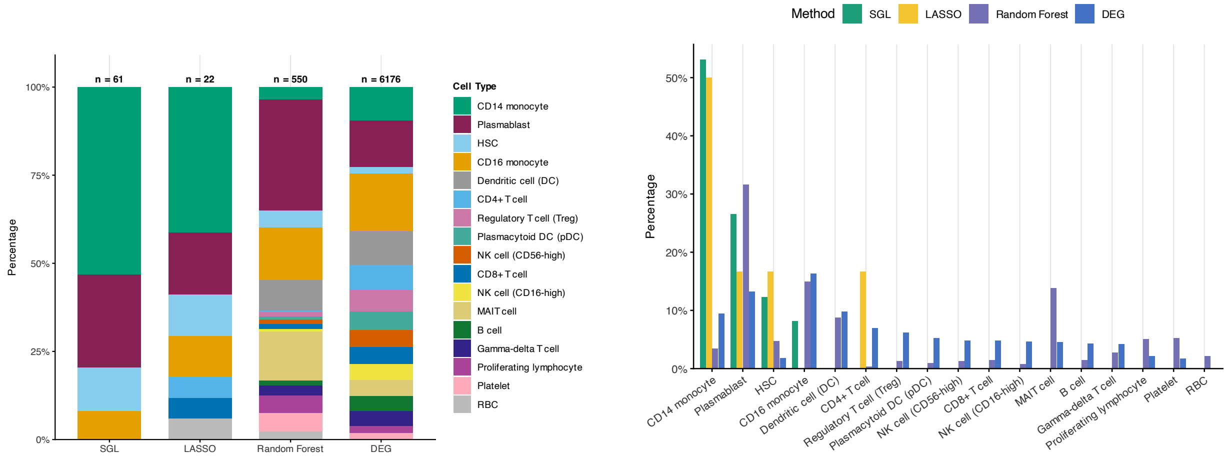

Figure S-13: Cell type composition plots of important features for the COVID dataset comparing SGL, Lasso, Random Forest, and Differential Gene Analysis. Random forest importance score filtered at 60. SGL and Lasso coefficient values filtered at 0.02. DEG adjusted p-values filtered at 0.05.

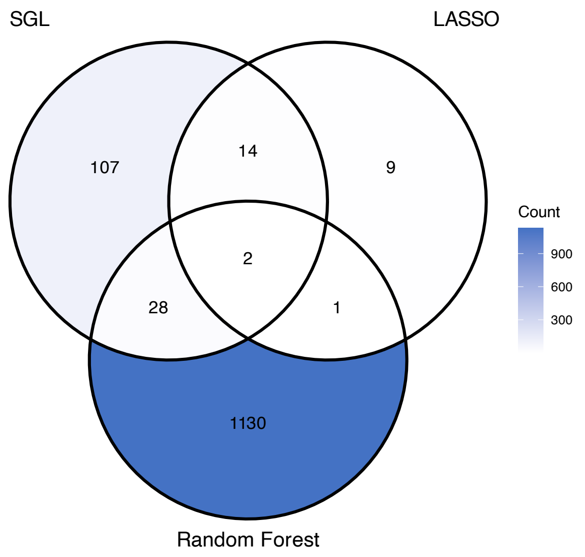


Figure S-14: Feature overlap across methods for COVID-19

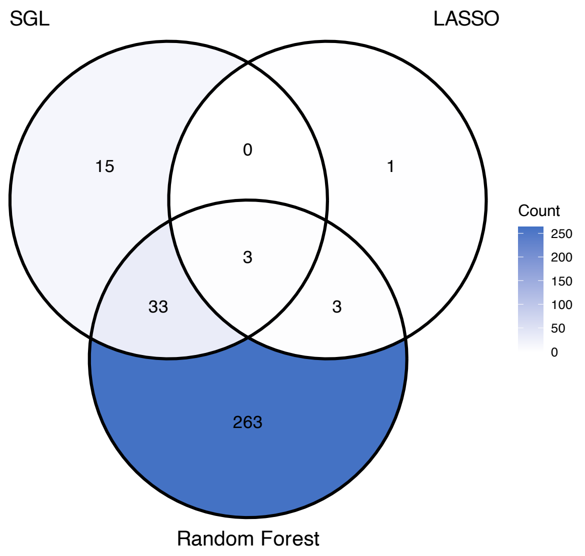


Figure S-15: Feature overlap across methods for CRC

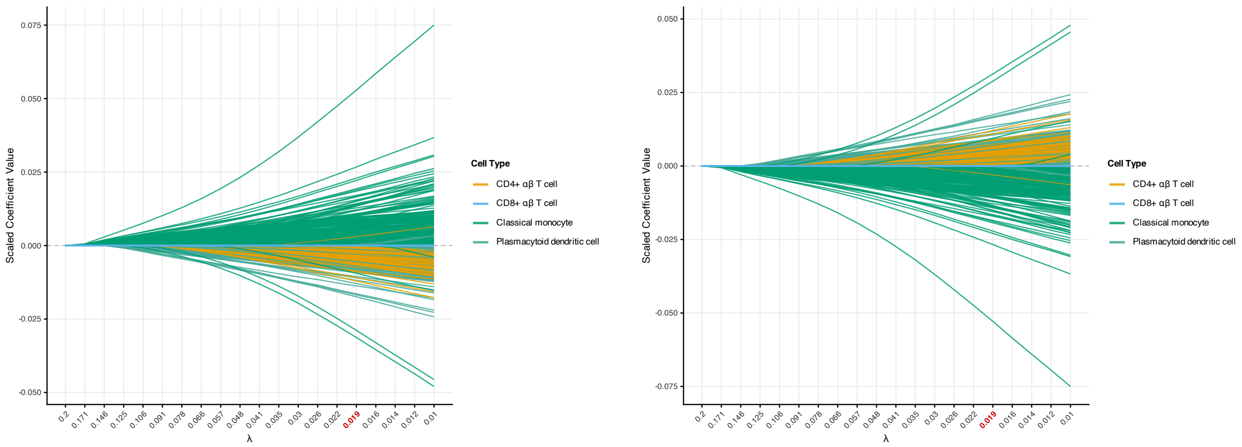


Figure S-16: Coefficient paths for the SLE dataset comparing managed condition (left) and normal condition (right) using SGL model trained on batch 2 only. No filtering on the coefficients values. The red tick mark indicates the CV-selected $\lambda$.

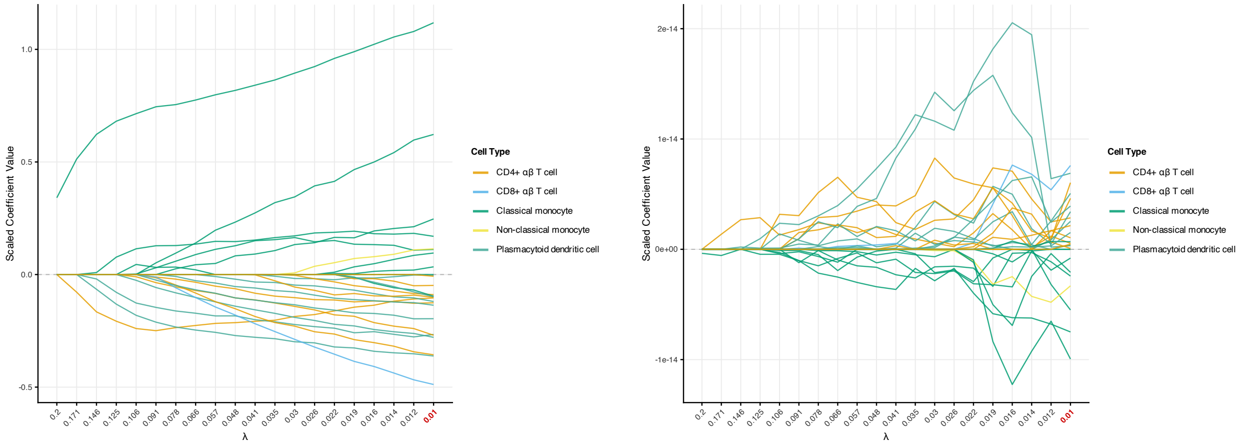


Figure S-17: Coefficient paths for the SLE dataset comparing managed condition (left) and normal condition (right) using lasso model trained on batch 2 only. No filtering on the coefficients values. The red tick mark indicates the CV-selected $\lambda$.

| **Cell Type** | **Number of DEGs** |
|---|---:|
| CD4-positive, alpha-beta T cell | 1,340 |
| Classical monocyte | 1,275 |
| Natural killer cell | 1,240 |
| Non-classical monocyte | 1,232 |
| CD8-positive, alpha-beta T cell | 1,197 |
| B cell | 1,139 |
| Conventional dendritic cell | 1,133 |
| Plasmacytoid dendritic cell | 900 |
| Lymphocyte | 686 |
| Plasmablast | 28 |

Table 4: Number of differentially expressed genes (DEGs) identified for each cell type in the SLE dataset (FDR $< 0.05$).

### S2 Supplementary Tables

| Cell Type | Number of DEGs |
|---|---|
| CD16 monocyte | 1,009 |
| Plasmablast | 819 |
| Dendritic cell (DC) | 603 |
| CD14 monocyte | 585 |
| CD4+ T cell | 430 |
| Regulatory T cell (Treg) | 381 |
| Plasmacytoid DC (pDC) | 321 |
| NK cells (CD56-high) | 298 |
| CD8+ T cell | 294 |
| NK cells (CD16-high) | 288 |
| MAIT cell | 280 |
| B cell | 263 |
| Gamma-delta T cell (gdT) | 257 |
| Proliferating lymphocyte | 131 |
| Hematopoietic stem cell (HSC) | 109 |
| Platelet | 108 |
| RBC | 0 |

Table 5: Number of differentially expressed genes (DEGs) identified for each cell type in the COVID-19 dataset (FDR $< 0.05$).

| Cell Type | Number of DEGs |
|---|---|
| Epi | 815 |
| TNKILC | 500 |
| Myeloid | 218 |
| Mast | 8 |
| Strom | 5 |
| B | 2 |
| Plasma | 1 |

Table 6: Number of differentially expressed genes (DEGs) identified for each cell type in the CRC dataset (FDR $< 0.05$).

| Cell Type | AUC |
|---|---|
| B cell | 0.582 |
| $CD8^+$ $\alpha\beta$ T cell | 0.578 |
| Classical monocyte | 0.576 |
| Conventional dendritic cell | 0.565 |
| Plasmacytoid dendritic cell | 0.559 |
| Non-classical monocyte | 0.554 |
| Progenitor cell | 0.533 |
| Lymphocyte | 0.517 |
| Natural killer cell | 0.507 |
| $CD4^+$ $\alpha\beta$ T cell | 0.505 |
| Plasmablast | 0.503 |

Table 7: Augur cell type prioritization results for the SLE dataset.

| **Cell Type** | **AUC** |
|---|---|
| $CD16^+$ monocyte | 0.792 |
| Plasmablast | 0.757 |
| $CD14^+$ monocyte | 0.678 |
| Dendritic cell | 0.670 |
| Platelet | 0.655 |
| $CD8^+$ T cell | 0.645 |
| Hematopoietic stem cell (HSC) | 0.622 |
| NK cell (CD16-high) | 0.612 |
| MAIT cell | 0.610 |
| $\gamma\delta$ T cell | 0.607 |
| Plasmacytoid dendritic cell (pDC) | 0.606 |
| B cell | 0.601 |
| NK cell (CD56-high) | 0.596 |
| $CD4^+$ T cell | 0.590 |
| Proliferating lymphocyte | 0.568 |
| Regulatory T cell (Treg) | 0.550 |
| Red blood cell (RBC) | 0.548 |

Table 8: Augur cell type prioritization results for the COVID-19 dataset.

| **Cell Type** | **AUC** |
|---|---|
| Epithelial | 0.589 |
| Plasma | 0.557 |
| Myeloid | 0.550 |
| Stromal | 0.538 |
| T/NK/ILC | 0.520 |
| B cell | 0.518 |
| Mast | 0.515 |

Table 9: Augur cell type prioritization results for the CRC dataset.
